# Dual GRIN lens two-photon endoscopy for high-speed volumetric and deep brain imaging

**DOI:** 10.1101/2020.09.19.304675

**Authors:** Yu-Feng Chien, Jyun-Yi Lin, Po-Ting Yeh, Kuo-Jen Hsu, Yu-Hsuan Tsai, Shih-Kuo Chen, Shi-Wei Chu

## Abstract

Studying neural connections and activities *in vivo* is fundamental to understanding brain functions. Given the cm-size brain and three-dimensional neural circuit dynamics, deep-tissue, high-speed volumetric imaging is highly desirable for brain study. With sub-micrometer spatial resolution, intrinsic optical sectioning, and deep-tissue penetration capability, two-photon microscopy (2PM) has found a niche in neuroscience. However, current 2PM typically relies on slow axial scan for volumetric imaging, and the maximal penetration depth is only about 1 mm. Here, we demonstrate that by integrating two gradient-index (GRIN) lenses into 2PM, both penetration depth and volume-imaging rate can be significantly improved. Specifically, an 8-mm long GRIN lens allows imaging relay through a whole mouse brain, while a tunable acoustic gradient-index (TAG) lens provides sub-second volume rate via 100 kHz ∼ 1 MHz axial scan. This technique enables the study of calcium dynamics in cm-deep brain regions with sub-cellular and sub-second spatiotemporal resolution, paving the way for interrogating deep-brain functional connectome.

## 1. Introduction

Brain is arguably the most complex organ, consisting of three-dimensional (3D) neural circuits that may span millimeter to centimeter with continuous spiking activities. However, even after more than 100 years of study since the era of Golgi and Cajal, our understanding toward the function of brain circuits is still far from complete, particularly due to the lack of a suitable tool that allows us to study neural connections and activities *in vivo* with 3D subcellular resolution in any location of a brain [1]. The challenges include not only penetration depth but also imaging speed of neural networks. The brain size of mammals is typically centimeter in scale, and in the mouse brain, ∼95% of neurons are not contained in the superficial cortical area [2]. In addition, millions of neurons interconnect and form complicated 3D networks, whose electrical and chemical signal dynamics are on the order of milliseconds to seconds. Therefore, it is highly desirable to develop brain imaging technologies with centimeter deep penetration and high-speed volumetric imaging capability.

For thick-tissue cellular observation in neuroscience, two-photon microscopy (2PM) has become a mainstream imaging technique [3]. There are several advantages of 2PM. First, it allows visualization of neuronal structure with sub-micrometer spatial resolution. Second, with its nonlinear nature, it provides intrinsic optical sectioning ability, which offers high contrast observation throughout a thick sample. Third, 2PM typically utilizes near-infrared light, which is less scattering and causes less damage than visible light in bio-tissue, thus enabling deep *in vivo* imaging. Unfortunately, due to strong scattering of brain tissues, typical penetration depth of 2PM in a mouse brain is less than 1 mm [4], which is inadequate for subcortical structures.

To enable observation of deep brain regions with light microscopy, several techniques have been developed. One strategy is to boost excitation power and the number of ballistic photons by integrating a regenerative amplifier; hence achieving ∼1 mm deep tissue imaging in the neocortex [5]. Another method is to utilize three-photon signals at long wavelength (1300 nm or 1700 nm), which is less scattered by tissue; thus, penetration depth can be improved to 1.2 mm [6,7]. Recently, a high-power, long-wavelength laser source can further overcome the depth limitation and achieve 1.5 mm deep imaging [8]. However, 1∼1.5 mm penetration depth is still far from penetrating through a whole mouse brain. In order to reach an arbitrary region of interest in a mouse brain, mechanical invasiveness is necessary.

Currently, endoscopy is one of the most practical methods of deep tissue imaging with minimal invasiveness [9]. By inserting a narrow, rod-like gradient-index (GRIN) lens, functions of neural circuits at several mm depth can be studied, by either single-photon [10,11] or two-photon endoscopy [12]. Although the former allows high-speed imaging of neural dynamics [13], the lack of optical sectioning and the vulnerability to scattering hinder its application in 3D imaging of neural circuits distributed across multiple layers. On the other hand, two-photon endoscopy [13–19] provides inherent optical sectioning, and is suitable for 3D imaging of thick brain regions. An emerging trend is to develop volumetric two-photon endoscopy. However, previously published two-photon volumetric endoscopy suffer from either low axial scanning speed [16,17] or low axial contrast [18,19], which cannot distinguish fast responses from two or more neurons that are overlapped in axial dimension.

Here, we demonstrate that by integrating another GRIN lens, i.e. tunable acoustic GRIN (TAG) lens [20–23] into two-photon endoscopy, high axial scanning speed with high axial contrast is achieved. Our dual GRIN lens two-photon endoscopy system allows *in vivo* imaging of neural circuits in centimeter-depth brain regions with high-contrast and sub-second volume rate, and has great potential for studying transient responses of 3D neural circuits with millisecond temporal resolution [24].

## 2. Materials and methods

### 2.1 Microscope setup

The high-speed volumetric two-photon endoscopic imaging system consists of a home-built two-photon microscope, a TAG lens, and a GRIN lens, as shown in Fig. 1. A tunable femtosecond-pulsed Ti:Sapphire laser (Chameleon Vision II, Coherent) was used as a light source for two-photon excitation. The laser beam was guided through a beam expander lens pair (LA1433-B & LA1131-B, Thorlabs) to match the numerical aperture of GRIN lens, and then was relayed onto galvo scanners (6215H, Cambridge Technology) to achieved raster scan. The raster scan pattern was relayed to an objective through a scan lens (SL50-CLS2, Thorlabs) and a tube lens (TL200-CLS2, Thorlabs). A dichroic beamsplitter (FF573-Di01-25×36, Semrock) was placed between the scan lens and the tube lens to separate the excitation beam and the epi-collected fluorescence photons. After the tube lens, a TAG lens (TAG lens 2.5β, TAG Optics) was placed directly on top of an objective (UPLFLN 10XP, Olympus) to enable high-speed axial scan of the focus. A 8-mm long GRIN lens (NEM-100-25-10-860-DS, Grintech), which was surgically implanted into a mouse brain (see section 2.4), was aligned with the objective. The scanned laser beam was relayed by the GRIN lens in deep mouse brain to excite two-photon fluorescence, which was then epi-collected by the same GRIN lens, objective, TAG lens, the tube lens, and the dichroic beamsplitter to a photomultiplier tube (PMT) module (H7422A-40, Hamamatsu). An image lens (LA1401-A, Thorlabs) was placed before the PMT to further converge fluorescence signals within the PMT detection area. A bandpass filter (FF01-520/15-25, Semrock) was placed between the image lens and the PMT to block residual excitation photons.

**Fig. 1.**
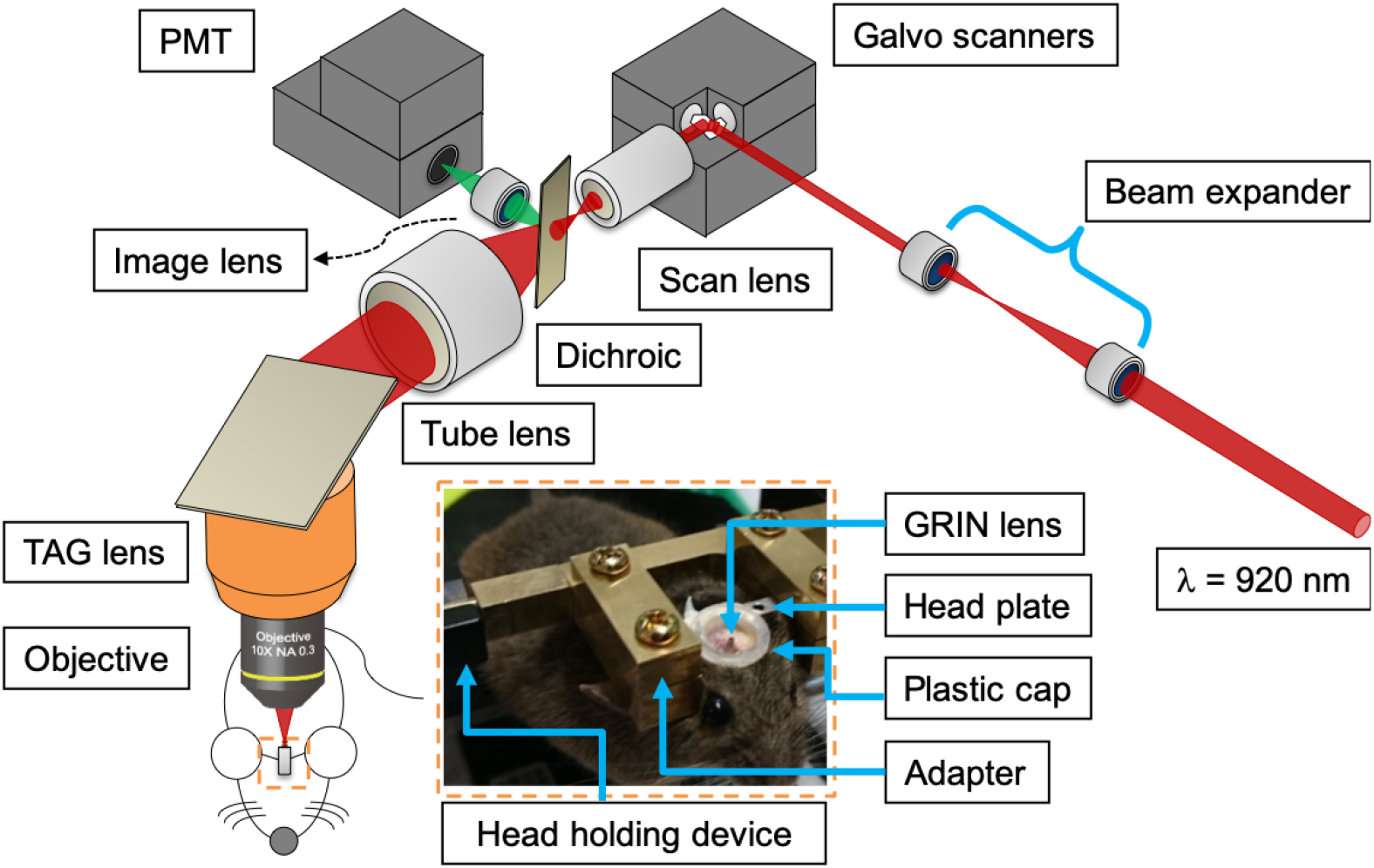
Dual GRIN lens two-photon endoscopic imaging system. Inset: A mouse implanted with a rod-like GRIN lens was head-fixed via a head plate and an adapter to the head holding device during the imaging session.

### 2.2 Fluorescent microspheres sample preparation

Fluorescent microspheres were prepared as a preliminary test sample. To characterize performance of GRIN lens, including its spatial resolution, extended depth of field (DOF), field of view (FOV), 500-nm diameter yellow-green fluorescent microspheres (F8813, Thermo Fisher Scientific, diluted 10^4^ times) dispersed into 0.4% agarose hydrogel were imaged. To mimic 3D observation of neurons, 10-μm diameter fluorescent microspheres (F8836, Thermo Fisher Scientific, no dilution) were used.

### 2.3 Mouse sample preparation

A three-month-old female heterozygous Ai96(RCL-GCaMP6s, Jackson Lab #028866) transgenic mouse was used for the experiments. The GCaMP6s gene was activated by cre recombinase vector that was delivered by adeno-associated virus 2.9 (AAV2.9, produced by Techcomm, National Taiwan University). All animal experiments were conducted according to the guidelines for the care and use of laboratory animals approved by the Institutional Animal Care and Use Committees of National Taiwan University. In the following, surgery sessions, including viral injection and GRIN lens insertion, and imaging session are stated sequentially.

In order to perform viral injection and GRIN lens insertion, the mouse was first mounted onto a stereotactic surgery platform. The mouse was fully anesthetized during all parts of the surgery. The anesthesia was induced by inhalation of 5% isoflurane for 2 min and maintained with 1-1.5% isoflurane for the rest duration of the surgery, with 1 and 0.2 L/min flow rate, respectively. The dosage of anesthetic followed guidance of the NIH Anesthesia and Analgesia Formulary. The mouse head was then shaved, and a 1-cm long incision was made at the scalp. The skin was then removed and the skull was exposed. A 1.5-mm hole was drilled through skull for GRIN lens insertion and two 0.8 mm holes were drilled at the lateral part of both parietal bones for two anchor screws. 300 nl of AAV2.9-cre virus was injected into the right suprachiasmatic nucleus (SCN) in 10 min (0.46 mm behind the bregma, 0.2 mm rightward to the midline, 6.0 mm beneath to the surface of skull). The flowing rate was controlled by a pump (KDS-310-PLUS, KD Scientific) and a 5 μl Hamilton 700 series glass syringe. After viral injection, GRIN lens insertion followed. On the stereotaxic apparatus, a customized holder was adapted to grab GRIN lens tightly, and the coordinate was set for the SCN. The GRIN lens was then gradually inserted into the mouse brain at a speed about 1 cm/hr to prevent tissue compression. After bleeding was staunched, a customized stainless-steel head plate (22 mm length, 3 mm width, and 1 mm thickness) was installed on the skull. Dental acrylic was applied to cement the GRIN lens, protecting ring and head plate over the whole skull surface and the two anchor screws. The ring was covered with a homemade plastic cap to protect the GRIN lens from damage and the mouse from contamination. The mouse was then accommodated alone for at least two weeks to recover.

During the imaging session, the mouse was first anesthetized by intraperitoneal injection of ketamine (Ketalar® Injection 50 mg/ml, Pfizer) and xylazine (Rompun® 20, Bayer) with 100 mg/kg and 135 mg/kg dosage, respectively. The dosage of anesthetic followed the guidance of the NIH Anesthesia and Analgesia Formulary. After the animal was fully anesthetized, the protection cap was removed and the GRIN lens was cleaned by 99.5% ethanol and air blowing. The mouse was then fixed onto a mouse head holding device (MAG-2, Narishige) via the head plate and a customized adapter. The mouse head holding device was screwed onto another translational stage (M-562-XYZ, Newport), to achieve tilt and 3D spatial alignment with a microscope objective.

### 2.4 Volume image reconstruction and data streaming

Two-photon fluorescence signals detected by the PMT were streamed and processed into volumetric images through the following steps. First, the PMT output was amplified with 50dB current-to-voltage gain (C6438-01, Hamamatsu). Second, the amplified voltage signals were converted into digital signals through a digitizer (NI-5734, National Instruments). Third, eight pulses were averaged in an FPGA module (PXIe-7972R, National Instruments) to form one voxel with enhanced signal-to-noise ratio (SNR). Last, voxels information was streamed to a solid-state disk (HDD-8261, National Instruments).

In order to reconstruct volumetric images from these voxels, data sampling with TAG lens and galvo scanners must be synchronized. In our system, as shown in Fig. 2, the femtosecond laser pulse train (80 MHz) was adopted as the external data sampling clock for the digitizer. Meanwhile, the TAG lens was sinusoidally driven at 70 kHz, and a synchronized TTL trigger was sent to the galvo scanner as a pixel trigger, i.e pixel dwelling time was ensured to be exactly the same as one period of TAG oscillation. Between two TTL triggers, the laser focus scan axially twice to collect ∼1,140 data points (80 MHz sampling), and were averaged and resampled to 73 voxels. As for galvo scanning, in our case, typically the line trigger is set to contain 128 pixels, and a frame trigger is set to contain 128 lines. As a result, a volumetric image with dimension 128 × 128 × 73 could be reconstructed with the aid of a custom MATLAB program.

**Fig. 2.**
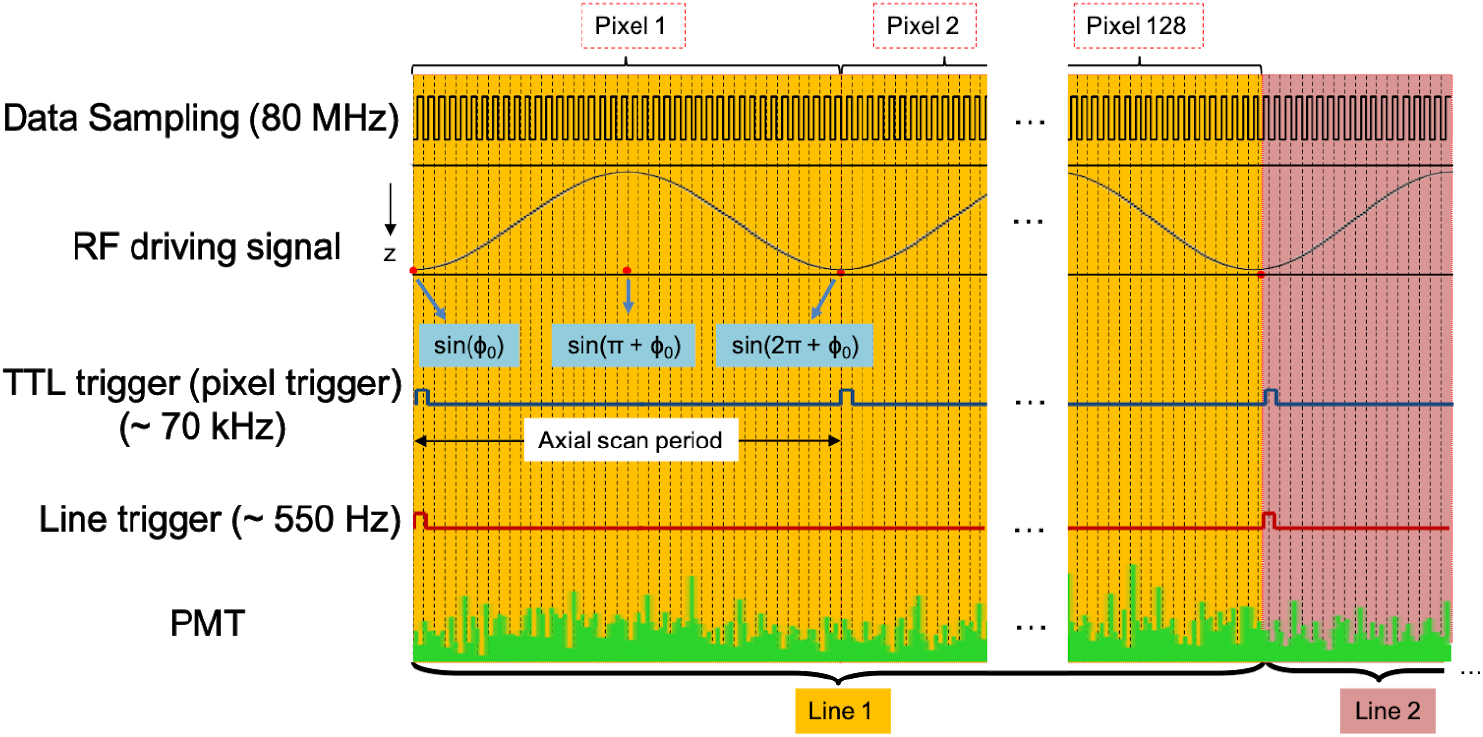
Signal flow chart for volume images reconstruction with synchronized data sampling, TTL trigger of the TAG lens, and triggers for galvo scanners.

## 3. Results

### 3.1 Spatial resolution of GRIN lens

Fig. 3 (a) and (b) show images of 500-nm fluorescent beads whose intensity profiles (averaged over 8 beads) along lateral and axial directions are presented respectively. The images were acquired near the center of GRIN lens FOV, and were normalized by maximum intensity of each bead. The red curves are Gaussian fittings of the two intensity profiles, and they reveal that full width at half maximum (FWHM), which characterizes spatial resolution, along lateral and axial directions are 1.03±0.06 µm and 10.64±0.70 µm respectively.

**Fig. 3.**
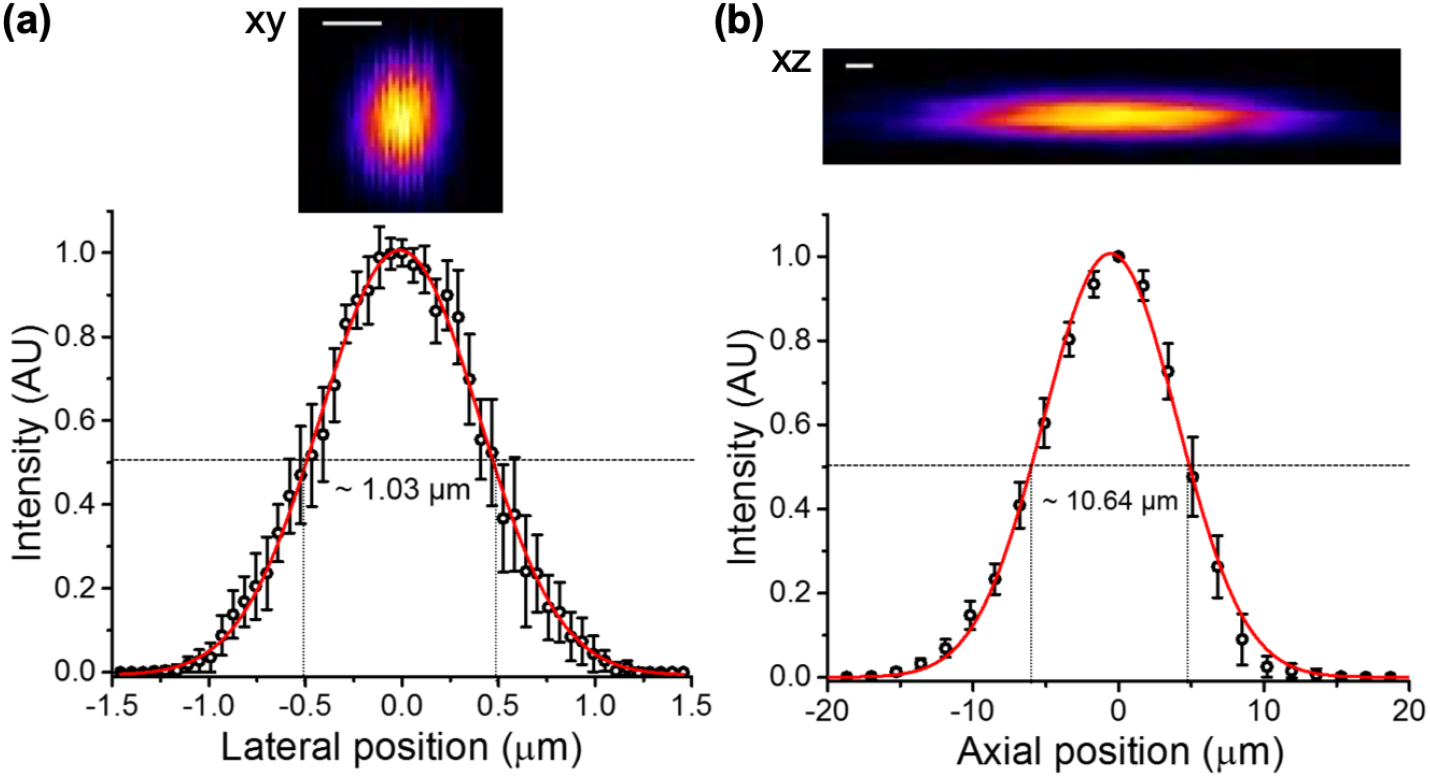
Lateral and axial point spread functions of the doublet, 1-mm-diameter GRIN lens obtained by imaging 500-nm-diameter fluorescent beads. (a) and (b) are representative PSF with average intensity profiles along lateral and axial direction, respectively. Scale bar: 1 µm.

### 3.2 Extended DOF measurements

The measurement of extended DOF was similar to that of axial resolution except that TAG lens was turned on. When TAG lens was turned on, the excitation laser focal spot was scanned over the z-axis at each pixel. Therefore, TAG lens resulted in an extended DOF on a pixel-by-pixel basis [21,22], and the extended axial intensity profile was adopted as the length of extended DOF according to a previous work [23].

Fig. 4 shows measurement of extended DOF, with an xz image and corresponding axial intensity profile and FOV. The incident beam size before TAG lens is 7 mm; hence, the resulting phase profile of TAG can be considered parabolic [22]. The red curve shows a dumbbell-like intensity profile, with its length ∼165 μm, which also determines the thickness of volumetric image stacks taken by our system will be. The xy intensity profiles at an endpoint (green), the center (yellow), and the other endpoint (blue) section of the extended DOF are also presented, showing lateral FWHMs are 1.09, 1.47, and 1.22 μm, respectively. The lateral FWHMs becomes slightly worse, which may result from additional aberration induced by TAG lens. However, the lateral FWHMs throughout the extended DOF still provide that system spatial resolution is good enough for most cellular imaging applications.

**Fig. 4.**
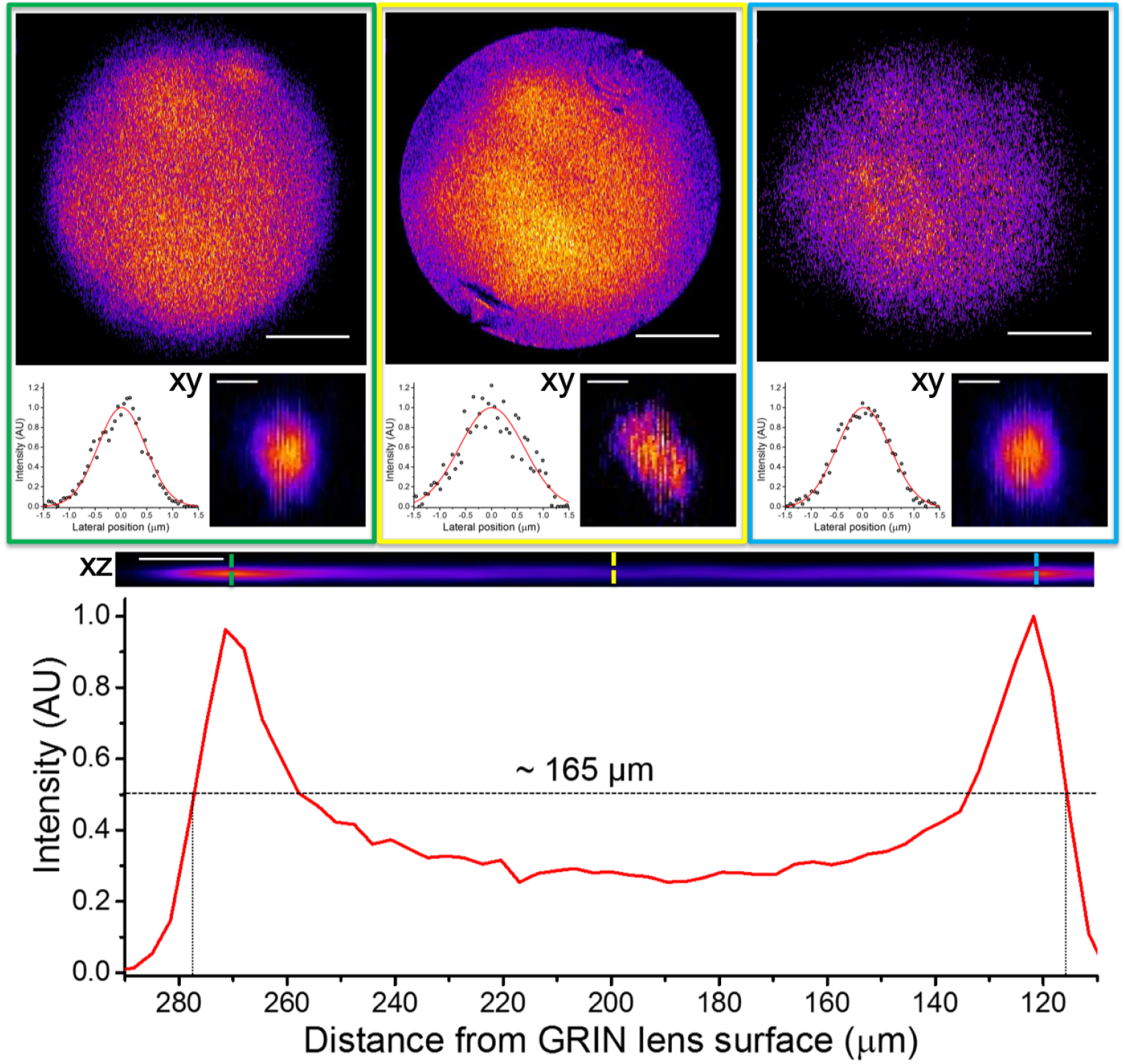
Extended DOF measured by using 500-nm fluorescent beads. Axial intensity profile with corresponding xz image is shown, with xy images and corresponding lateral intensity profiles and FOV at different sections. Scale bar: 10 μm for xz image, 1 μm for xy images, and 100 μm for FOV images.

According to literatures, GRIN lens FOV at different focal depth may be limited by off-axis aberrations [25,26]. Therefore, GRIN lens FOV was also measured to characterize the size of imaging volume. The measurement of GRIN lens FOV was similar to that of lateral resolution except that galvo scanners were set to scan across as large areas as possible. During the measurement, the relative position between the GRIN lens and the beads sample was kept constant and the distance between the objective and the GRIN lens was varied for the GRIN lens to focus at different focal depths. Then FOVs at different depths were acquired according to proposed criteria [16,19].

As shown in the upper-most three panel of Fig. 4, the size of FOV was ∼ 350 μm in diameter and remained almost the same at different focal depths. Compared with theoretical value (380 μm), our measured FOV is ∼ 13% smaller. Although adaptive optics are proposed to improve, current FOV at different focal depths is still large enough to cover many deep brain nuclei such as SCN (size ∼ around 200 × 200 × 200 μm^3^) [27].

### 3.3 Applications of volumetric endoscopic imaging

After characterizing the resolution and imaging volume of view, here we demonstrate two representative applications of applying the high-speed volumetric endoscopic system to monitor dynamics of biological tissues. One is to capture flow speed of fluorescent microspheres that mimicked blood flow. The other one is to achieve *in vivo* functional imaging of neuronal activity from SCN, which is located at the bottom side of a mouse brain.

#### 3.3.1 Phantom of blood flow via fluorescent microspheres

Fig. 5 shows time-lapsed series of 10-µm fluorescent microspheres flowing to the right inside a tube, mimicking blood cell flow or a cellular microfluidic system (see Visualization 1 for full video). These images demonstrate that our temporal resolution achieves ∼4-Hz volume rate, with spatial resolution more than enough to resolve individual cells. The high spatiotemporal resolution allows us to determine the flow speed to be ∼ 15 µm/s.

**Fig. 5.**
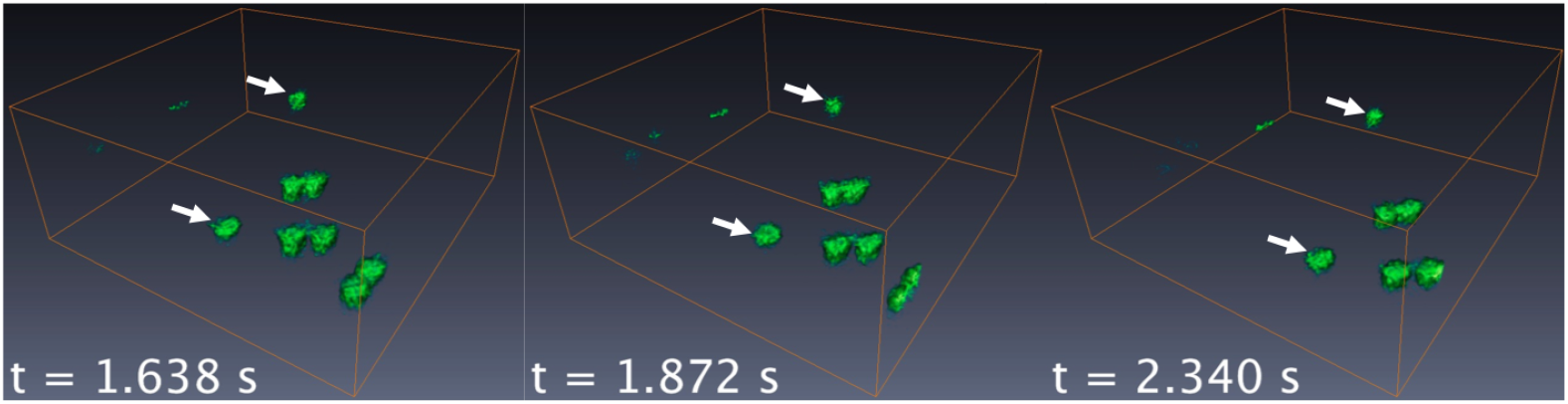
Representative time-lapsed images of flowing 10-µm fluorescent microspheres in 3D. The white arrows in each panel indicate the flowing direction of microspheres. Volume size: length ∼ 180 µm, width ∼ 180 µm, and height ∼ 165 µm.

#### 3.3.2 Volumetric functional imaging of neurons in living mice deep brain regions

Fig. 6 shows *in vivo* functional imaging of spontaneous neuronal activities from SCN of a head-fixed anesthetized mouse. Fig. 6(a) is a volumetric image of GCaMP6-labeled SCN (see Visualization 2 for time-lapsed functional response). The volume size is approaching 200 × 200 × 200 µm^3^, which is enough to cover most of the SCN in one brain hemisphere of the mouse. Within the volume, six regions of interest (ROIs) were manually selected and indicated as the white circles. The calcium transients (ΔF/F) of these regions are shown in Fig. 6(b). ROI 1 and 2 show burst-like neural responses and responses of ROI 3 and ROI 4 gradually increase in time. ROI 5 reveals slight decay of calcium signal and ROI 6 shows almost no response. The diverse patterns indicate that the observed fluorescence changes reflected neural responses instead of motion artifacts.

**Fig. 6.**
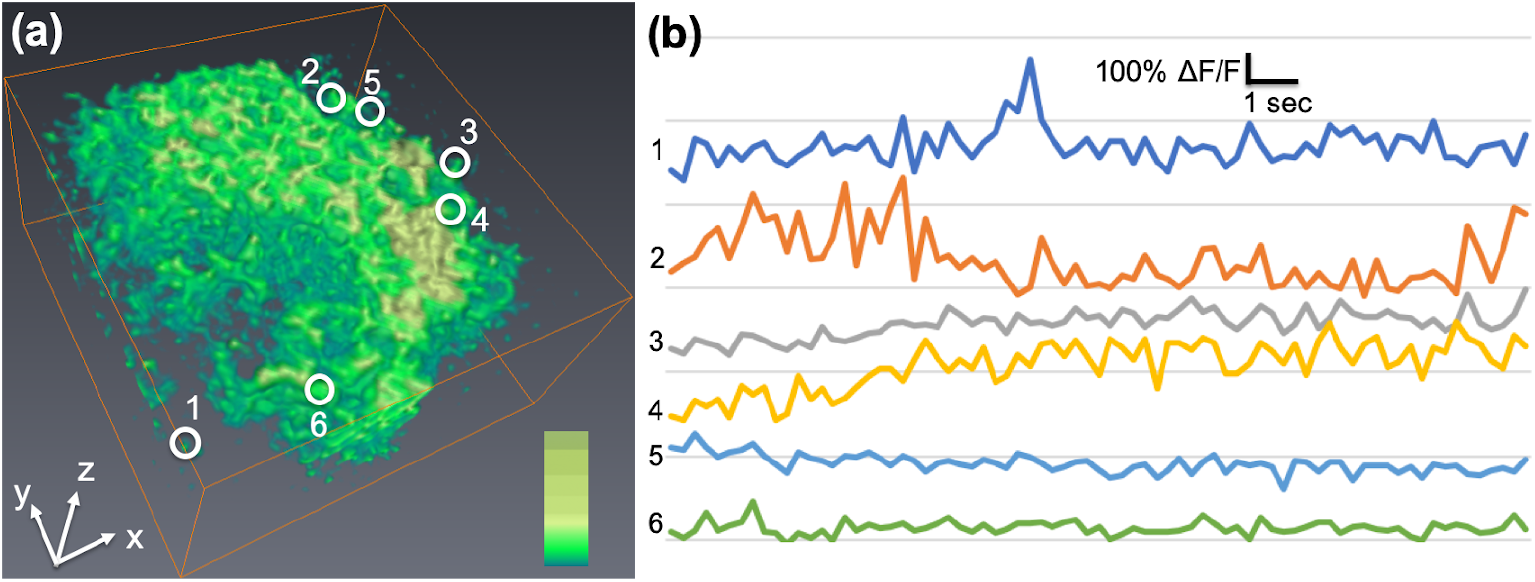
*In vivo* functional imaging of neuronal activity from SCN of a head-fixed anesthetized mouse. (a) 3D reconstruction of a SCN with 0.25-sec temporal resolution. The volume size is ∼ 180 × 180 ×165 µm^3^. (b) shows neuronal activity measured by the calcium transient ΔF/F in those circular ROIs of (a).

## 4. Discussion

In Fig. 3, we have shown that the lateral and axial resolution of our endoscopy system are ∼1 µm and ∼10 µm, respectively. Compared with theoretical values of a NA 0.5 GRIN lens, our measured FWHM in lateral and axial directions are respectively 51.5% and 79.1% larger. In table 1, spatial resolution of recent endoscopy literature based on a NA 0.5 GRIN lens is listed. Surprisingly, most of the other results are also much larger than theoretical prediction. The poor resolution of GRIN lens may result from spherical aberration [16,28] or birefringence [29], and adaptive optics [16] or GRIN lens cascade [29] were proposed to improve spatial resolution.

**Table 1.** Comparison of GRIN lens spatial resolution

| Type of NA 0.5 GRIN lens | Lateral resolution | Axial resolution |
| --- | --- | --- |
| Theoretical value | 0.693 $\mu\text{m}$ | 5.92 $\mu\text{m}$ |
| NEM-100-25-10-860-DS, current GRIN lens | 1.03±0.06 $\mu\text{m}$ | 10.64±0.70 $\mu\text{m}$ |
| Same GRIN lens in [19] | 1.23 $\mu\text{m}$ | 12.1 $\mu\text{m}$ |
| NEM-050-25-10-860-DM, in [16] | 1.29 $\mu\text{m}$ | 12.4 $\mu\text{m}$ |
| NEM-050-25-10-860-S, in [18] | 0.81±0.04 $\mu\text{m}$ | 8.55±0.35 $\mu\text{m}$ |
| SLW, in [17] | 2.3 $\mu\text{m}$ | 62.5 $\mu\text{m}$ |

In Fig. 6, we have demonstrated *in vivo* volumetric functional images in SCN (6-mm deep in a mouse brain). In fact, Our dual GRIN lens two-photon endoscopy may have other applications. For brain study, since SCN lies almost at the bottom of a mouse brain, most deep brain regions, such as hippocampus, amygdala, and substantia nigra [16, 17] could also be examined and studied using our system. Apart from brain study, most biological systems are intrinsically dynamic, such as muscle contraction [30] and blood flow [31]. In Fig. 5, we have demonstrated flowing fluorescent beads mimicking blood cells flow. Based on the largest volume size (∼ 350 × 350 ×165 µm^3^) and temporal resolution (∼4-Hz volume rate) of our system, the maximum flowing speed can be estimated is ∼1500 µm/s. In fact, Li. *et al*. [32] estimated flowing speed of red blood cells in mice to be ∼600 µm/s under comparable FOV and temporal resolution, which means our technique is definitely able to measure blood flow speed in deep brain regions.

## 5. Conclusions

In conclusion, we have demonstrated a novel two-photon volumetric endoscopic imaging system, which was accomplished by integrating a GRIN lens and a TAG lens into a home-built two-photon microscope. Such a novel technique allows *in vivo* imaging of neurons in a centimeter-depth mouse brain with high-contrast and sub-second volume imaging rate.

## 5.2 Acknowledgments

We appreciate Dr. Tsai-Wen Chen for providing the design of the mouse head plate.

